# Clustering Electrophysiological Predisposition to Binge Drinking: An Unsupervised Machine Learning analysis

**DOI:** 10.1101/2024.04.24.590901

**Authors:** Marcos Uceta, Alberto del Cerro-León, Danylyna Shpakivska-Bilán, Luis M. García-Moreno, Fernando Maestú, Luis Fernando Antón-Toro

## Abstract

**Background:** The demand for fresh strategies to analyze intricate multidimensional data in neuroscience is increasingly evident. One of the most complex events during our neurodevelopment is adolescence, where our nervous system suffers constant changes, not only in neuroanatomical traits, but also in neurophysiological components. One of the most impactful factors we deal with during this time is our environment, especially when encountering external factors such as social behaviors or substance consumption. Binge Drinking (BD) has emerged as an extended pattern of alcohol consumption in teenagers, not only affecting their future lifestyle, but changing their neurodevelopment. Recent studies have changed their scope into finding predisposition factors that may lead adolescents into this kind of patterns of consumption.

**Methods:** In this article, using unsupervised machine learning (UML) algorithms, we analyze the relationship between electrophysiological activity of healthy teenagers and the levels of consumption they had two years later. We used hierarchical agglomerative UML techniques based on Ward’s minimum variance criterion to clusterize relations between power spectrum and functional connectivity and alcohol consumption, based on similarity in their correlations, in frequency bands from theta to gamma.

**Results:** We found that all frequency bands studied had a pattern of clusterization based on anatomical regions of interest related to neurodevelopment and cognitive and behavioral aspects of addiction, highlighting the dorsolateral and medial prefrontal, the sensorimotor, the medial posterior and the occipital cortices. All this patterns, of great cohesion and coherence, showed an abnormal electrophysiological activity, representing a dysregulation in the development of core resting-state networks. The clusters found maintained not only plausibility in nature, but robustness, making this a great example of the usage of UML in the analysis of electrophysiological activity, a new perspective into analysis that, while contributing to classical statistics, can clarify new characteristics of the variables of interest.

## Introduction

As the data load of neurosciences keeps growing, working with its complexity and dimensionality becomes more difficult. This is why the need for original perspectives of analysis is becoming more pressing. In this regard, the emergence of innovative technologies, such as machine learning (ML), has proven to be a valuable alternative to traditional statistical analyses **[1,2,3]**.

Originating in the mid-20^th^ century, ML appeared in pursuit of predicting new outcomes, using intrinsic characteristics from the data where the algorithm trained, to generalize and classify unseen data **[4]**. This, the supervised ML (SML), may be the most commonly known usage of ML. Furthermore, we can find unsupervised ML (UML), a set of techniques that allows us to analyse raw data without pre-existing labels, not predicting any outcome, but grouping data by its similarities or reducing dimensionality of the general dataset by eliminating redundancy. The application of UML has increased as novel technologies have been refined, influencing several lines of research, like disease identification **[5, 6, 7]** and differential analyses of traits of interest in healthy population **[8, 9, 10]**.

The UML approaches include hierarchical agglomeration algorithms, which involve methods that group variables within the data based on similarities of their characteristics. This framework cluster variables in groups, from one (where all data is enclosed) to *k* groups (where each variable is its own group). These approaches allow us to discretize dataset in a *k* number of groups with an optimal trade-off between information and coherence in order to achieve better explainability **[11]**.

Numerous criteria have been designed to group variables based on their similarity. When aiming to cluster complex data (such as neuroimaging and electrophysiology data), it is crucial to minimize internal noise within the clusters. Ward’s minimum variance criterion **[12]**, an enhanced version of the Lance-Williams dissimilarity formula, facilitates the coherent grouping of data improving compactness of clustering and maximizing the information captured within them **[13]**. This technique facilitates extracting relevant information from complex interactions.

As noted by Sadaghiani and colleagues **[14]**, electrophysiology and, mostly, brain connectivity contain a high degree of complexity, especially when the system is affected by abnormal situations, like a disease or a misregulation. Moreover, brain’s complex dynamics are extremely sensible to both internal and external factors which may contribute to alter its functioning and subsequent behavioural outcomes. In this scenario, UML emerge as a useful approach to disentangle the complex associations between brain’s functioning data and behavioural profiles.

Adolescence is a critical developmental stage where brain dynamics change towards a more organized system. During this period, the brain is undergoing a general maturation across several functional systems, radically changing its electrophysiology, with important consequences on cognition and behavior. One of the most common lifestyle factors emerging during adolescence is risk-taking behaviors, and more concerning, intensive alcohol consumption or Binge Drinking (BD). Binge Drinking (BD) **[15]** is characterized by the ingestion of high doses of alcohol (at least 4 standard alcohol units (SAUs) for women and 5 for men) within shorts periods of time **[16]**. This pattern of consumption has been demonstrated to be harmful to the adolescent’s brain, due to its acute vulnerability against external factors during its development **[17]**. More precisely, BD has been shown to lead to abnormal structural **[18,19]**, functional **[20, 21, 22]** and neuropsychological impairments **[23]**, which can have enduring consequences for individuals.

However, a major milestone in the study of adolescent BD it’s the unravelling of predisposition profiles that lead some individuals to engage in BD years later, a field of study of increasing relevance **[24]**. Some neuroanatomical studies have identified brain’s structural differences, including a reduction in the average volume of certain prefrontal, temporal and parietal regions before BD onset **[25, 26]**. Regarding neurofunctional evidence, fMRI studies have reported decreased BOLD signal among prefrontal and parietal regions **[27, 28]**, tightly associated with behavioural control circuits. Additionally, electrophysiological studies using magnetoencephalography (MEG) have found increased functional connectivity profiles associated with future BD patterns, both in inhibitory control networks **[29, 30]** and resting-state networks **[31]**. All these findings suggest neurodevelopmental abnormalities linked to the development of alcohol misuse. Nevertheless, despite the growing body of evidence concerning this matter, the specific relationship between complex electrophysiological profiles and future BD behaviours remains poorly understood. After a thorough inspection of the literature associated with the key words characteristic of the work proposed in this study (“electrophysiology” or “MEG” or “EEG”, “predisposition” or “risk”, “binge drinking” and “adolescence”), we could see that the number of published articles is small, and even smaller if we add the component of the use of new analysis tools such as ML.

With that objective, this prospective study applies UML techniques in the identification of electrophysiological patterns associated with future alcohol BD in teenagers. We used a database containing resting-state MEG signals from adolescents before the onset of alcohol consumption, and their relationship with consumption habits two years later. Using Ward’s minimum variance clustering, we explored the interaction between functional connectivity and power spectra profiles in association with future alcohol consumption. This framework allowed us to identify patterns of electrophysiological activity, distinctive of higher alcohol intake years later and topographically consistent with the organization of several function system and its development.

## Methods

### Participants

The sample for this study was comprised of adolescents recruited from various schools in the Region of Madrid, as part of two independent longitudinal projects funded by the Spanish *Ministerio de Sanidad* in 2015 and 2017. Both projects followed the same assessment protocol, consisting of two evaluation stages separated by a two-year follow-up period. Prior to the experiment, all participants reported no history of alcohol consumption or familiar alcohol use disorder. Additionally, all subjects successfully completed the alcohol use disorder identification test (AUDIT) **[32]**, and individuals who reported prior alcohol use were excluded from the study. In the first stage (before alcohol use onset), 148 adolescents agreed to participate in the first arm of the study, undergoing a semi-structured interview about their drug use habits to exclude any potential consumers; later, they underwent a magnetoencephalography (MEG) study, consisting in five-minute eyes-closed resting-state, from which 142 also agreed to participate in an MRI study.

After a follow-up period of two years, in the second stage, 104 of these participants underwent the AUDIT test and a semi-structured interview again, aiming to measure their alcohol consumption habits. Using this information, we calculated the quantity of standard alcohol units (SAUs) consumed during regular drinking episodes for each participant, considering the number of beverages consumed within a 2–3-hour period. Tobacco and cannabis use were controlled by excluding those participants with regular use of these substances. Finally, after quality control of the MEG and MRI data, a final sample of 103 subjects (mean age 13.75 ± 0.64, 53 females) completed the entire protocol and were selected for analysis.

The sample consisted almost entirely of native-born participants and was balanced in terms of weight and height; elements such as parental education, participants’ auto-informed quality of life or dedication to the study were also controlled for (for a more detailed explanation of demographics, see **Supplementary Table 1**). Informed consent was obtained from all participants and their parents or legal guardians during the first stage of the study, following the guidelines outlined in the Declaration of Helsinki. The ethical committee of *Universidad Complutense de Madrid* granted approval for the study.

### MEG recordings

The recordings were conducted at the Cognitive and Computational Neuroscience Laboratory (UCM-UPM) of the Biomedical Technology Centre (CTB) (Madrid, Spain) using an Elekta Neuromag system with 306 channels (Elekta AB, Stockholm, Sweden) placed inside a magnetically shielded room (VacuumSchmelze GmbH, Hanau, Germany). We acquired 5 minutes of task-free (resting-state) data with eyes closed using an online bandpass filter between 0.1 and 330 Hz, and a sampling rate of 1000 Hz. To later calculate their electrical components and subtract them from the brain signal, we also acquired eye blinks and heartbeat using two sets of bipolar channels with the same configuration. To aid in the source reconstruction stage, the head shape of the participants was also obtained using a Fastrak three-dimensional digitizer (Polhemus, Colchester, Vermont).

We initially preprocessed the data using a spatial filtering provided by the manufacturer (tSSS) **[33]** with MaxFilter software (v 2.2, Elekta AB, Sweden), tuning the parameters to 0.9 as correlation limit and 10 seconds as length of the correlation window. Then, we used the FieldTrip packages **[34]** to detect artifacts and remove interfering signals (noise, heart beats, ocular activity) with a SOBI-based ICA **[35]**. Finally, we segmented the data into 4-second epochs, avoiding the artifacted segments.

### Source Reconstruction

As source model, we defined a homogeneous grid with 1 cm of separation between sources using the MNI template, yielding 2459 source positions inside the cranial cavity. These sources positions were labelled using the AAL atlas **[36]**, and only those 1206 positions designated as one of the 78 cortical areas were considered for further analysis. With the help of a linear normalization between the MNI space standard T1 image and the subject specific T1 image, we transformed this source model into a subject space, segmenting the image into the different tissues using the unified segmentation algorithm in SPM12 **[37]**. This combination was used to build a realistic single shell interface representing the inner skull cavity. Then, we transformed this source and volume conduction model into the MEG space using the head shape as aid.

We solved the forward problem by building a lead field based on a modified spherical solution **[38]**, and the inverse problem by using a LCMV beamformer **[39]**, finally recreating the source-level activity. The spatial filter was build using LCMV and the covariance matrix generated from the broadband (2 to 45 Hz) sensor-space activity, filtered using a 1800^th^-order FIR filter. To avoid any possible distortion, data was filtered in two passes, and using 2 seconds (2000 samples) of real data at each side of the epoch as padding.

### Power Spectrum

We estimated the relative source-level activity for specific frequencies of each band (theta (4-8 Hz), alpha (8-12 Hz), beta (12-20 Hz), and gamma (30-45)) using a *mtmfft* method with *dpss* as windowing function, with a 1 Hz smoothing. This data served as the precise dataset of brain activation in each of the AAL 78 cortical regions for each frequency band.

### Functional Connectivity

We used the phase locking value (PLV), which examines consistency of the phase differences between two time-series, to evaluate functional connectivity (FC). To determine the PLV **[40, 41]** for each frequency and epoch, we used the Hilbert transformation to extract the instantaneous phase of each signal node at each specified time, and then estimated the synchrony between each pair of signals using the difference of their phases. Both the band-pass and the Hilbert filters were applied over the epochs including 2 seconds of real data as padding (removed before continuing the analysis). We obtained the whole-brain FC matrix, estimating the PLV between each pair of cortical sources position. Then, we estimated the PLV between each pair of the 78 cortical regions using the root mean square of the PLV values. Finally, we calculated the FC strength of each ROI as the average PLV value of that ROI.

### Unsupervised Machine Learning Workflow

Although the usage of independent variables (FC values, power spectrum and alcohol consumption) was plausible, our work focused on the relationship between these variables, not the variables themselves. To this aim, we decided to work with Spearman correlation matrices between the electrophysiological variables and the alcohol consumption, as the relationships between variables are expected to be non-linear.

We followed a pipeline (**Figure 1**) using predominantly Python libraries. Firstly, to determine the optimal number of groups into which our dataset could be divided, we assessed the cluster quality with varying numbers of groups. This involved using *k-means* automatically grouping of the original dataset, iterating with a range of groups from 2 (the minimal number of groups possible) to 10 (a quantity estimated as large enough to be adequate). The efficiency of the groups produced was then assessed by computing the within-cluster sum of squares (WCSS), as well as cohesion measures such as Silhouette score **[42]**, the Calinski-Harabasz index **[43]**, and the Davies-Bouldin index **[44]**. Consequently, the optimal number of groups was determined for the dataset, considering it not as the strictly best number but rather as the one yielding the highest consistency.

**Figure 1:**
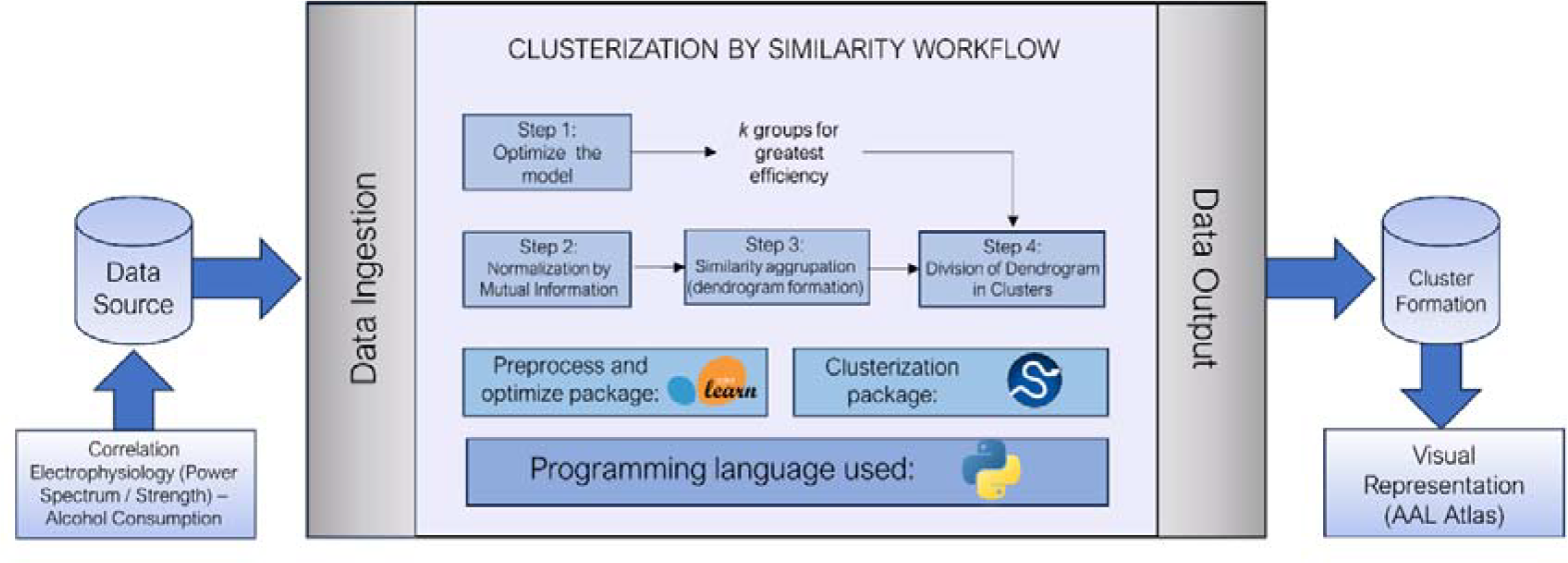
Schematic overview of the general workflow, representing the initial data sources (Spearman correlations between power spectrum/functional connectivity and alcohol consumption), the general steps within the pipeline, with the packages and the programming language used for them; and the data output, with the visual representation. inscribed in the AAL atlas, of the clusters formed.

We performed this analysis using the tools provided by the Sci-kit learn metrics package (https://scikit-learn.org/stable/modules/classes.html#module-sklearn.metrics). All metric values for the range of groups *k* described above are depicted at large in **Supplementary Table 2**.

To enhance the information contained within the working dataframe, as well as normalizing the data and reducing the internal noise, improving the coherence of the clustering, we added an additional step, performing a normalization of the data based on mutual information **[45]**. This technique, emerged from information theory, characterizes the amount of information from a variable *x* within a variable *y*, measuring the redundancy between variables, and resulting in a measure of the uncertainty of all our variables by eliminating this redundant information. The correction applied to the original dataset, then, is based on this measurement, specifically from the variation of information **[46]** within the system, a measure of the total entropy (or novel information) each step of the correction. Applying a correction of the data in each iteration, we filter the noise, increasing novel information content, thereby enhancing the system’s stability. The package utilized to calculate the mutual information score was sourced from the Sci-kit learn package mentioned before.

Finally, with this corrected dataset, we performed a data clustering by similarity using agglomerative hierarchical UML algorithms. We considered the previously obtained division value *k* for the group number, to direct the algorithm towards obtaining *k* separated clusters. We found, across all bands, that the most efficient number of groups our dendrogram should be cut was 2. We performed the same pipeline for several methods and, given its superior stability across various iterations of the model and its robustness in generating coherent groupings, we opted for the minimum variance model or the Ward’s method:

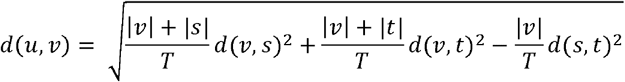

where u is the newly joined cluster consisting of clusters *s* and *t*, v is an unused cluster within the data frame, and T=|v|+|s|+|t| **[47, 48]**. We used Euclidean distance to quantify the distance between clusters (as a measure of dissimilarity between variables).

The ultimate representation and analysis of the data following the data clustering process was carried out using dendrogram visualizations, tree-like structures where data is grouped based on the distance between variables. For each frequency band, we obtained a dendrogram which identifies, within distinct groups, the cortical ROIs associated with each type of variable (strength-consumption or power spectrum-consumption).

All the methods utilized for unsupervised hierarchical agglomeration, as well as the representation in dendrograms, were sourced from the SciPy toolkit (https://docs.scipy.org/doc/scipy/index.html).

## Results

### Theta Band

Theta band clustering defined two separate groups with an evident and uneven separation (see **Fig. 2A**). Cluster 1 has an average intergroup distance of 1.4149 (SD = 0.8439), with a general cutoff at 4.6187. We found that negative Power-BD correlations appeared in this cluster in parietal cortex (somatosensorial cortex, supramarginal gyrus), as well as in the middle cingulate and temporal cortices. Specifically, the Heschl’s and the superior temporal gyri were included only for the right hemisphere. On the other hand, we found that positive Strength-BD variables appeared in the occipital regions (left lingual and inferior occipital gyri), as well as in the posterior cingulate cortex and superior parietal and angular gyri (see **Fig. 3A-C**).

**Figure 2:**
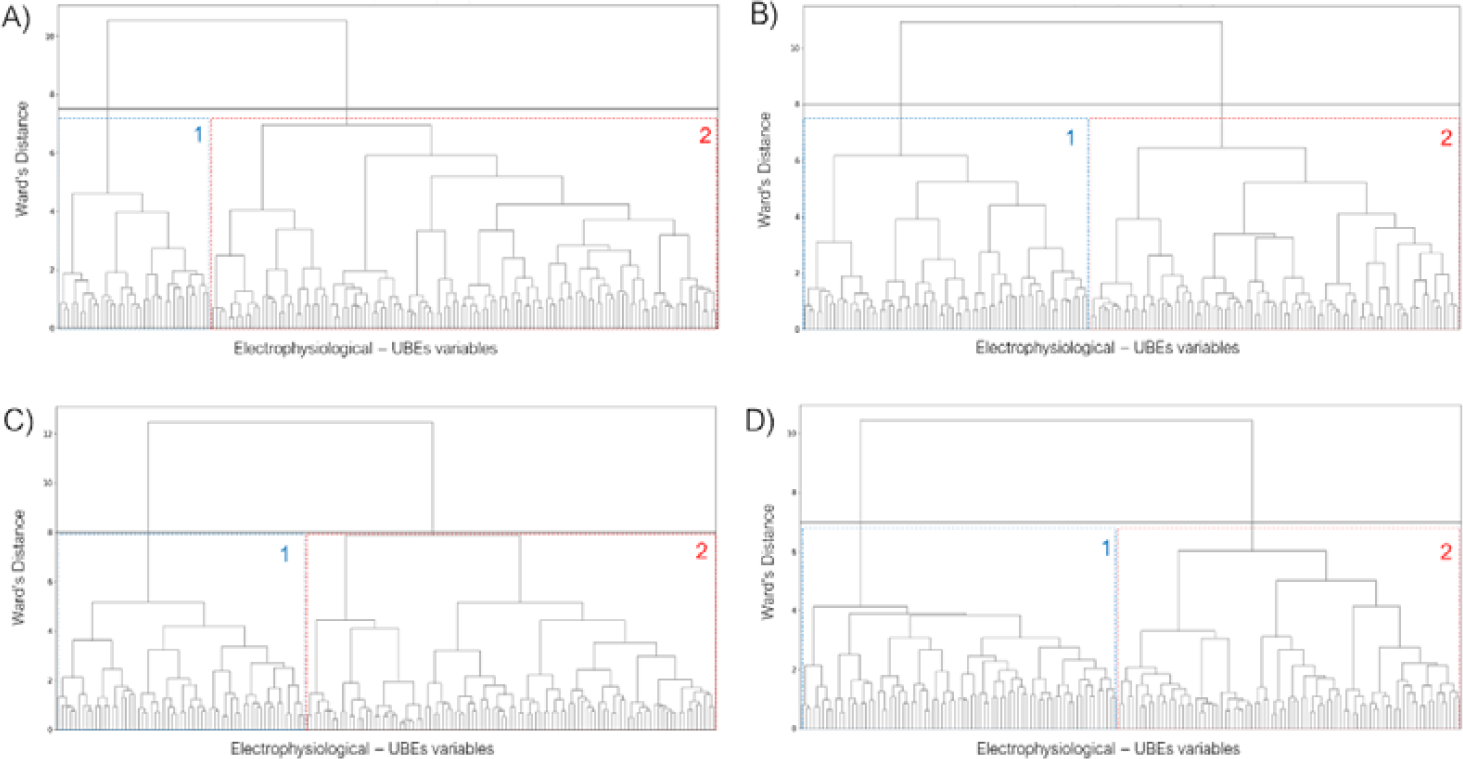
Dendrogram-based depiction of the distribution of correlations among electrophysiological variables (power spectrum and strength) and consumption (SAUs), generated through unsupervised machine learning, utilizing Ward’s minimum variance criterion, for each frequency band (**A**: theta. **B**: alpha. **C**: beta. **D**: gamma). For each band, within the overall grouping: 1 (in blue) = cluster 1 grouping: 2 (in red) = cluster 2 grouping.

**Figure 3:**
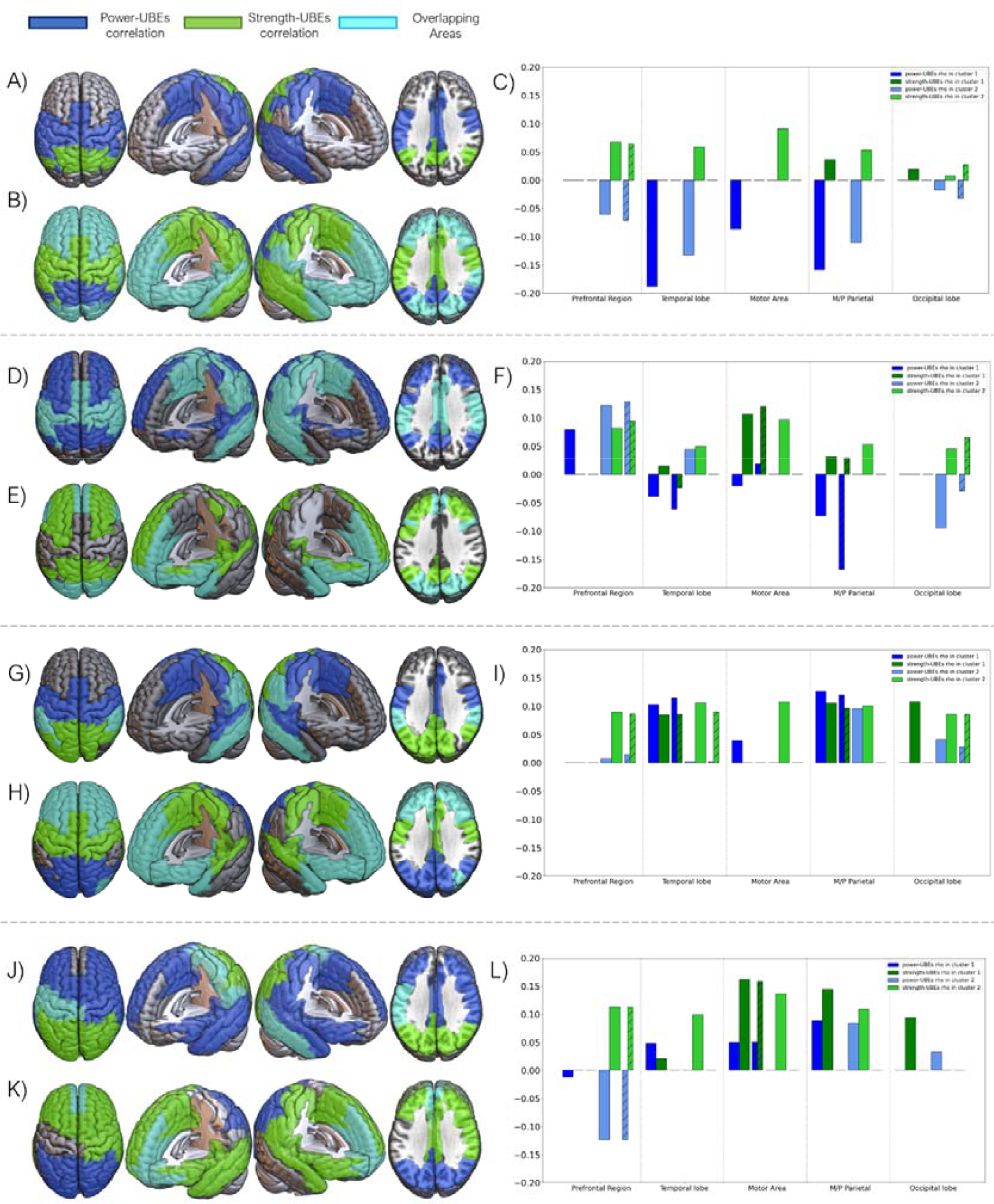
Visual representation of the grouping, into two distinct clusters, of correlations among electrophysiological variables (power spectrum and strength) and alcohol consumption (SAUs) for the four frequency bands of interest (theta, alpha, beta and gamma, in descending order). **A, D, G, J**: Visual representation of variables within cluster 1 for each frequency band. **B, E, H, K**: Visual representation of variables within cluster 2 for each frequency band. **C, F, I, L**: Correlation graph between electrophysiological variables and alcohol consumption for each lobe and the cingulate cortex, for both clusters within each frequency band. In a stripped pattern are the values of the overlapped ROIs within each group of areas.

The vast majority of the variables were found within Cluster 2. This cluster is characterized by an average intergroup distance of 1.3553 (SD = 1.0799), breaking away from the general group at 6.9489. Negative power-BD and positive strength-BD correlations overlapped, in this cluster, at the prefrontal regions, anterior cingulate cortex, superior temporal gyrus, and occipital lobe. Individually, negative power-BD correlations appeared at the angular gyrus, the posterior cingulate cortex and precuneus; on the other hand, positive strength-BD variables were found in the parietal cortex (somatosensorial cortex, supramarginal gyrus) as well as in the middle cingulate cortex and paracentral and fusiform gyri (see **Fig. 3B-C**).

For a detailed report of each individual region in both clusters, see **Supplementary Figure 1A**.

### Alpha Band

Clustering in alpha band represented an unbalanced division of variables (see **Fig. 2B)**. Cluster 1 has an average intergroup distance of 1.4360 (SD = 1.0560), with a general block cutoff at 6.1684. Negative power-BD correlations were found in parieto-temporal regions (including the precuneus, and the posterior paracentral and angular gyri), with the addiction of the positive power-BD correlations on the dorsal superior frontal gyrus. Positive strength-BD correlations appeared overlapping with power-BD variables in the supplementary motor area, middle cingulate cortex, parietal regions, and temporal areas (plus the hippocampus and parahippocampus) (see **Fig. 3D-F**).

Regarding cluster 2, it exhibits an average intergroup distance of 1.3237 (SD = 1.0491), with a general block cutoff occurring at 6.4650. Positive power-BD variables were shown around occipital regions, as well as the temporal poles and the anterior cingulate cortex; negative power-BD correlations were found in the frontal lobe. On the other hand, positive strength-BD variables appeared in the same areas but taking all frontal lobe (with the exception of the right precentral gyrus) as well as all occipital lobe and part of the temporal poles. Both variables overlapped at the orbitofrontal gyri, as well as in the temporal poles and the most posterior part of the occipital lobe (see **Fig.3E-F)**.

For a detailed report of each individual region in both clusters, see **Supplementary Figure 1B**.

### Beta Band

Clustering in beta band represent an uneven but distributed clusterization (see **Fig. 2C**). Cluster 1 was identified as a highly cohesive cluster, with an average distance of 1.4473 (SD = 0.9320), breaking away from the general block at 5.1751. Positive power-BD correlations appeared in parieto-temporal regions (all temporal gyri, as well as Heschl’s gyrus), and the middle cingulate cortex. On the other hand, positive strength-BD correlations were found with a more occipital orientation, reaching also the posterior cingulate cortex, as well as the middle and inferior temporal gyri. Both variables overlapped there, as well as in the supramarginal and the fusiform gyri (see **Fig. 3G-I)**.

Cluster 2 has an average intergroup distance of 1.2977 (SD = 1.1304), with a cutoff of the general group at 7.8791. Both positive power-BD and positive strength-BD correlations overlapped at the temporal poles, as well as in the frontal lobe (excepting for precentral gyrus, not present for Power-BD) and in the anterior cingulate cortex and insula. Positive power-BD also appeared in the occipital lobe, as well as in the posterior cingulate cortex. Positive strength-BD, on the other hand, appears in the parietal lobe (superior parietal and angular gyri, and precuneus) and the middle cingulate cortex (see **Fig. 3H-I**).

For a detailed report of each individual region in both clusters, see **Supplementary Figure 1C**.

### Gamma Band

Gamma band clustering defined an even and balanced division of variables, (see Fig. 2D). Cluster 1 exhibits an exceptionally high internal coherence, with an average distance of 1.4750 (SD = 0.7640), and an early separation of the general block at 4.1232. Positive power-BD correlations were shifted towards the right hemisphere, taking the entire frontal lobe (except for the middle and orbital gyri) as well as the temporal and the anterior part of the parietal lobe (paracentral, postcentral, and supramarginal gyri). Positive strength-BD correlations were shown at the occipital lobe completely. Both variables overlapped at the left precentral gyrus, as well as the left postcentral and supramarginal gyri, the paracentral gyrus, and the right middle temporal gyrus (see Fig. 3J-L).

Cluster 2 has an average intergroup distance of 1.3567 (SD = 0.9564), diverging from the general block at 6.0243. Positive power-BD correlations appeared in the occipital lobe, whereas negative power-BD correlations were found at orbitofrontal areas. On the other hand, positive strength-BD correlations are shifted towards the right hemisphere, appearing in the temporal lobes as well as in the frontal and right parietal regions (right postcentral and supramarginal gyri). Both variables overlapped at the anterior cingulate cortex, as well as in the orbitofrontal areas and the gyrus rectus (see **Fig. 3K-L)**.

For a detailed report of each individual region in both clusters, see **Supplementary Figure 1D**.

### General analysis of precision of clustering

In order to ascertain the viability and explainability of the clustering performed, we replicated the general pipeline in a random set of data. The results, shown in the **Supplementary Figure 2**, are a clear demonstration of a random patching of cerebral areas in both Power-BD and Strength-BD correlations, different from the results found in real electrophysiological data clustering. We also assessed the efficiency of different clustering criteria, choosing Ward’s minimum variance as the most plausible algorithm we could use. The different clustering made by the different criterion can be visualize in the **Supplementary Figure 3**.

## Discussion

The application of ML techniques has witnessed a progressive increase in recent years. Their growing renown can be attributed to their ability to discern previously undetected patterns within datasets, which can pave the way for novel hypotheses and subsequently inform future analytical processes. In the present study, UML methodologies were utilized to elucidate the correlations between electrophysiological variables, including power spectra and FC, and the subsequent emergence of alcohol consumption habits in adolescents. Through this methodology, we were able to discern unique clustering patterns, each exhibiting a uniform cortical distribution and displaying distinct associations with alcohol consumption across different frequency bands.

### Machine Learning Results

Most of the technical analysis in this paper is based on the examination of the cohesion and coherence between the different variables under investigation. Electrophysiological brain activity presents nonlinear relationships that, when computed in association with environmental traits (like alcohol intake), increases its complexity, making it crucial to provide a new perspective other than classical statistics. The type of data we encounter in the various frequency bands is inherently distinct, as was demonstrated by the clustering results depicted above. Depending on the frequency range we are dealing with, the achieved cohesion varies from the maximum homogeneity observed in the slowest band (theta), to the maximum heterogeneity observed in the division of the fastest, and possibly noisiest, band (gamma). These unequal responses not only speak to the nature of the different frequencies, but also to the precision of the algorithm’s division; even when analysing noisy frequency bands, the clustering depicted keep a consistency with ecological characteristics of the brain. In this sense, the richness in the nature of the different frequency bands does not reduce the explainability of the algorithm, finding plausible groups.

Applying rigorous clustering criteria such as the Ward’s minimum variance method has proven highly suitable for the information considered in this work. As we can observe when comparing the results obtained from random data to those obtained from actual data, the separation and grouping of variables within the anatomically defined areas of the brain is coherent, while also maintaining bilateral symmetry. Other less stringent (yet not simplistic) algorithms (such as single linkage), or methods based on geometric criterion (like centroid or median linkage), yield less precise groupings. In fact, ad hoc analyses were performed, finding that, as we can observe in **Supplementary Figure 3**, the clusterings found using these other different criteria are practically random; algorithms that apply weights to the clusterings (like weighted linkage) introduce smoothing in the divisions, subsequently losing information. It is necessary, therefore, to obtain groupings that not only consider the intragroup cohesion of each cluster, but also the comparative coherence of one cluster with the next; in this sense, the Ward’s algorithm provides both capabilities, rendering it ideal for our scenario **[13]**.

### Electrophysiological results

Overall, our findings indicate that power spectra and FC variables coalesce into two distinct similarity clusters, primarily characterized by four unique, anatomically driven patterns of activity: dorsolateral and medial prefrontal, sensorimotor and temporal, medial and posterior parietal, and occipital patterns. These patterns are uniformly distributed in the cortex across frequency bands, yet they exhibit diverse interactions amongst themselves. Notably, these patterns are in line with the disposition of resting-state networks (RSn) identified in electrophysiological MEG data by Brooke and colleagues **[49]**, clustering cortical activity in crucial nodes of the default mode network: the frontoparietal network, the sensorimotor network, the medial parietal network and the visual network. Concerning the relationship between power and strength variables with alcohol misuse, the distribution of these RSn within the two similarity clusters contrasts between each other; this is, regions pertinent to one variable in cluster 1 correspond to those related to the other variable in cluster 2. These results evidence the complex and nonlinear interactions between power and FC dynamics within cortical networks.

General results showed that higher FC across frequency bands and functional networks are related with higher future alcohol use. On the other hand, power exhibited diverse relationships depending on the frequency and the cortical network. Interestingly, slower frequency bands tend to exhibit an inverse relationship of power and strength variables with alcohol use. Contrary, within faster frequency bands both power and strength variables showed positive correlations with alcohol consumption. However, there are notable exceptions, concerning the prefrontal power within the alpha and gamma bands, which reversed these patterns. Such divergent patterns of prefrontal alpha and gamma bands are coincident with their unique distribution within the prefrontal cortex, separating the activity of medial and orbital regions from the dorsolateral part. Such regions have been demonstrated to be the core of the neuromaturative changes of the adolescence’s brain **[50]** and have been found to be particularly associated with the potential development of risk behaviours such as substance consumption **[50, 51, 52]**.

### Relevance and relationship with prior findings

Prior studies have pointed the neurological underpinnings predisposing adolescents to substance consumption, often highlighting the differences of neural pathways regulating self-control and risk assessment **[53, 54]**. This relationship gains complexity when considering prior research indicating variances in RSn among adolescents, which are thought to underlie various cognitive and behavioural aspects of addiction. Studies have shown alterations in RSn in adolescents with substance abuse disorders, including those prone to binge drinking **[20, 55, 56, 30]**. These networks are instrumental in self-referential thinking, emotional regulation, and cognitive control, respectively, and their dysregulation has been associated with increased impulsivity and risk-taking behaviours, prevalent in individuals with adolescent alcohol misuse **[57]**. Our observation of the distinct cortical activity clusters coincides with these studies, suggesting a distinct interplay between various cortical networks and their developmental trajectories. For instance, the heightened sensitivity in the dorsolateral and medial prefrontal regions, as corroborated by our power and FC dynamics, resonates with their established role in decision-making and emotional regulation, faculties often compromised in adolescents prone to binge drinking **[58]**. Furthermore, these cortical complexity works as a potential mediator of behaviour, reflecting the neuroadaptive processes, critical during adolescence **[50]**.

Research on the electrophysiological signatures of predisposition to BD is currently limited, and it is therefore challenging to connect present findings with existing literature. Nevertheless, the observed outcomes can be evaluated in the context of developmental trajectories. Established research has considered typical neurodevelopment of electrophysiological markers during adolescence **[59, 60]**. Broadly, throughout adolescence, there is a decline in power-spectra density in the theta band (in posterior regions), whilst there is an increase in beta and gamma band (in the latter, in the prefrontal regions), having a mixed evolution in alpha band (increase in temporo-parietal regions and reduction in prefrontal regions). Given this backdrop, our results suggest that a precocious maturation profile within specific cortical areas and frequency bands could correlate with increased future consumption tendencies. In contrast, patterns discerned in the alpha band for both prefrontal and parieto-occipital zones seem to present an inverse correlation with the conventional neuromaturational trajectory. This inverse correlation is also noticeable in the prefrontal pattern of power in gamma band, suggesting a contradictory link between consumption and cortical power compared to standard neurodevelopmental progression. In relation to FC, a consistent increase across all frequency bands is noted during adolescence, with the exception of the gamma band, which exhibits a reduction with age. Interestingly, our findings indicate that pronounced alcohol misuse correlates with elevated FC in the gamma band, especially within the frontal cortex. All these neuroadaptive mechanisms might identify individuals more susceptible to substance use as a function of both the unique neural maturational profiles and the concurrent psychosocial pressures typical of this developmental stage.

### Relevancy of machine learning, future implications, and limitations

The results obtained in this work underscore two key points regarding the utility of ML in the analysis of neurophysiological variables. Firstly, it prompts consideration for the optimization and refinement of analyses that would involve a substantial effort, performing complementary analyses to those made by classical statistics. The use of such tools has successfully navigated studies that would otherwise have been impossible or exceedingly laborious and costly **[61, 62, 63]**. This also mitigates potential biases that researchers may introduce when analysing or processing their data. Within the field of neuroscience, reducing noise and enhancing the coherence of analyses is important, and the application of these automated techniques might contribute to increase precision and objectivity **[64, 65, 66]**, giving support to the existing knowledge. Secondly, the study of neuroscience aims to go beyond the analysis of isolated variables, but rather the examination of interactions among them, with a holistic perspective. In this context, the application of non-linear methodologies that allow for the visualization or identification of links among variables is essential. Techniques such as those used in this work, or machine learning strategies like reinforcement learning (RL) **[67, 68]**, are invaluable. They significantly diminish the dimensionality and intricacy of the data, displaying the information into a comprehensible structure. These advanced approaches are crucial for refining complex datasets, particularly in studies dealing with multifaceted electrophysiological information and its behavioural outcomes.

The findings presented here are notably consistent with empirical observations in natural settings. The extraction of patterns could be used in the future as a foundational source of information for training convolutional neural networks (CNNs) that predict the occurrence of intensive alcohol consumption behaviours in adolescents solely using non-invasive neuroimaging measures. This would enable the identification of potential future consumers, allowing institutions to increase the resources provided to prevent these behaviours.

These findings, while pointing in a direction that aligns with prior knowledge and providing new insights into the next steps to be taken, have several limitations worth highlighting. The use of ML techniques, as employed in this article, poses significant issues regarding explainability. In neuroscience, the use of induction algorithms like these can be highly beneficial, but it is crucial to minimize reliance on the “black boxes” that machine learning algorithms rely on to function. Peering into these boxes to avoid being left with prediction values lacking relevant information can be costly and requires the use of simpler techniques or a thorough understanding of the methods employed. The use of UML or RL and deep RL techniques may be a step toward adding explainability, albeit at the expense of some efficiency and robustness compared to the more commonly used methods.

## Conclusion

This work has confirmed that areas closely related to the neurodevelopment of teenagers, such as the prefrontal cortex, the DMN, and the somatosensory cortex, are pivotal areas within the clustering performed by Unsupervised Machine Learning techniques, serving as essential patterns within the observed dynamics, and providing new information regarding abnormal neurodevelopment as predisposing factor towards alcohol consumption. These findings provide clear foundations for future studies in the search for predisposition patterns and the assessment of alcohol consumption in adolescents, as well as providing new insights on the usage of novel automatic techniques.

