## Supplementary figures and images for "Clustering Electrophysiological Predisposition to Binge Drinking: An Unsupervised Machine Learning analysis"

### Supplementary Figure 1

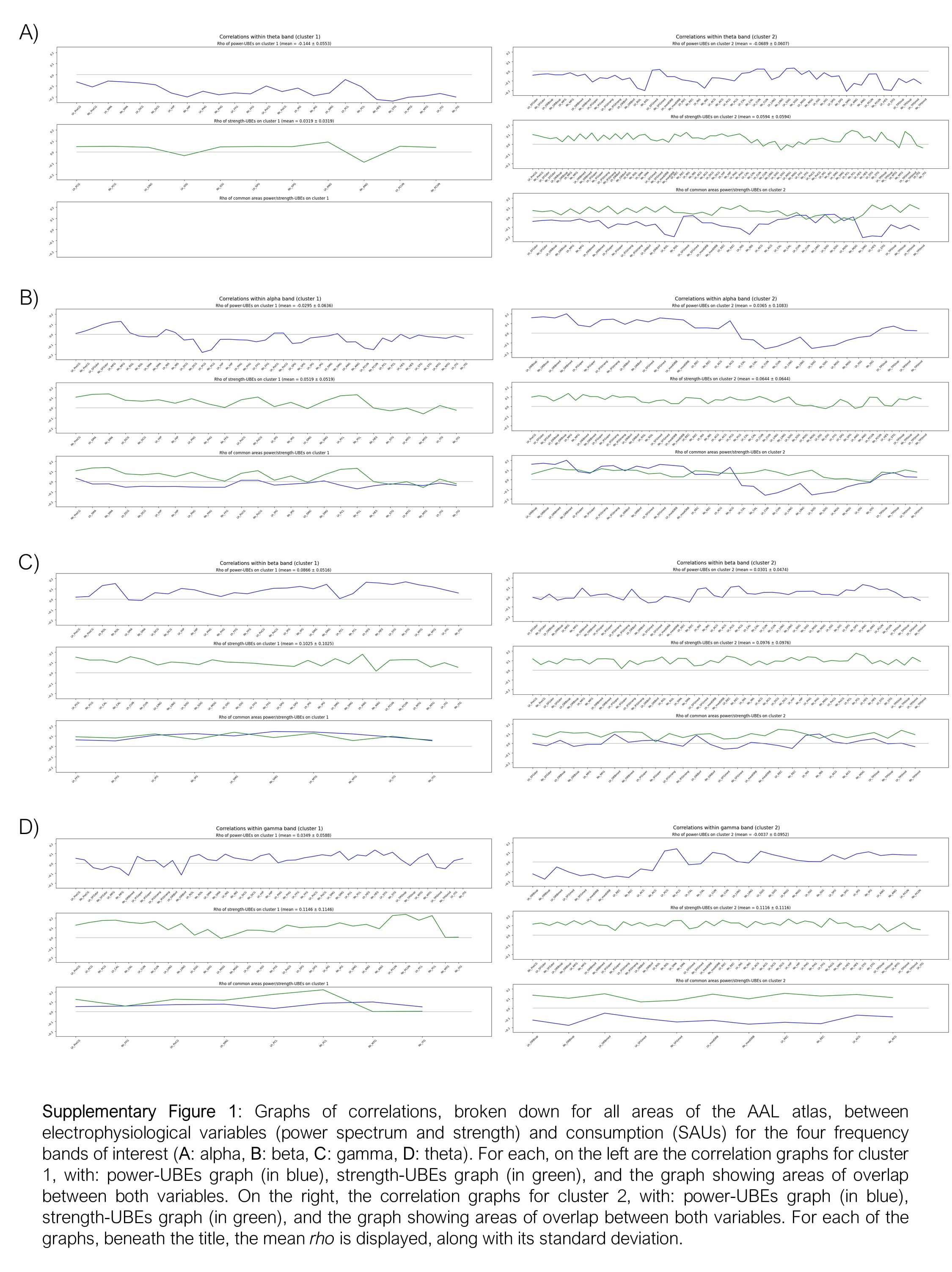

### Supplementary Figure 2

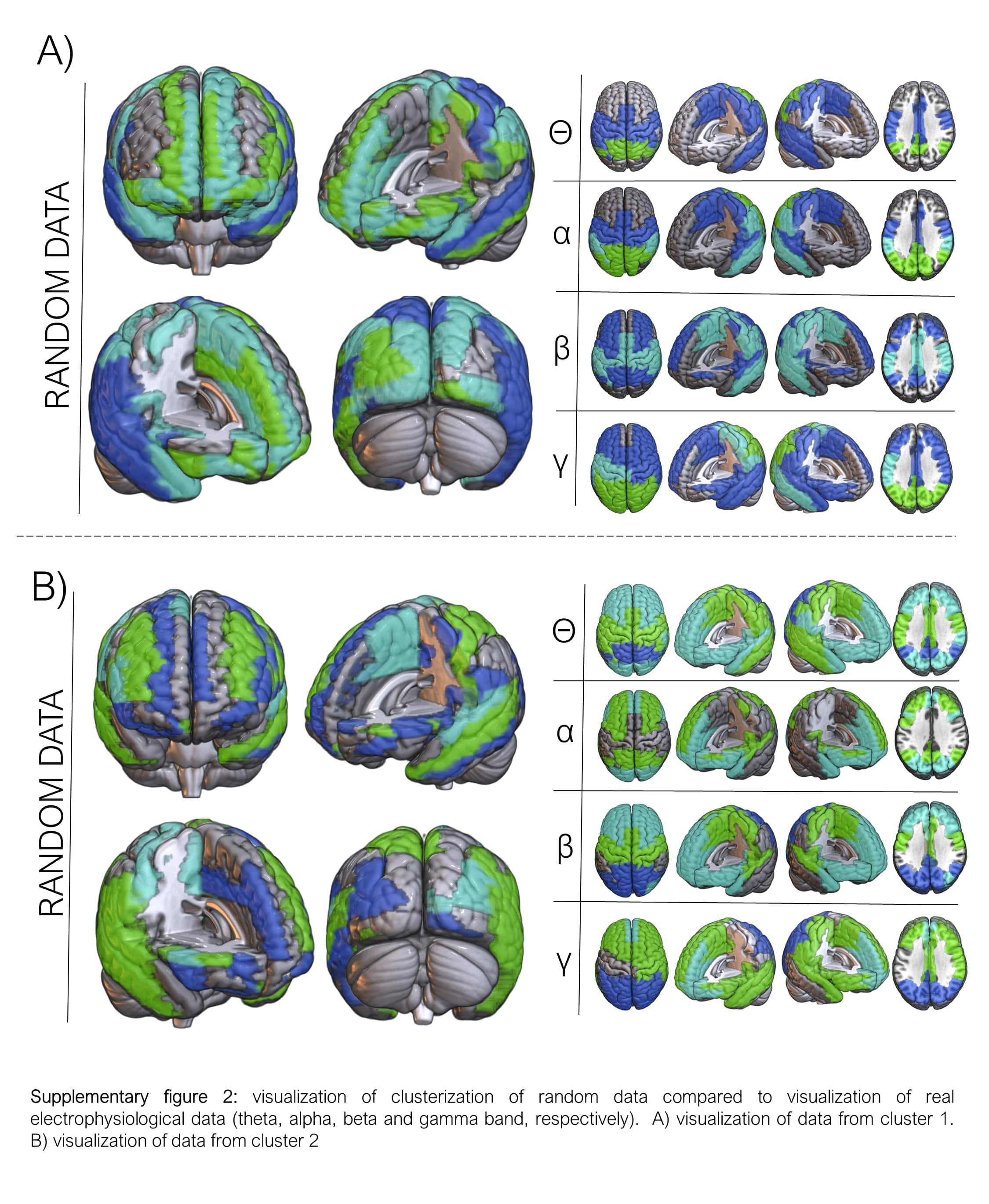

### Supplementary Figure 3

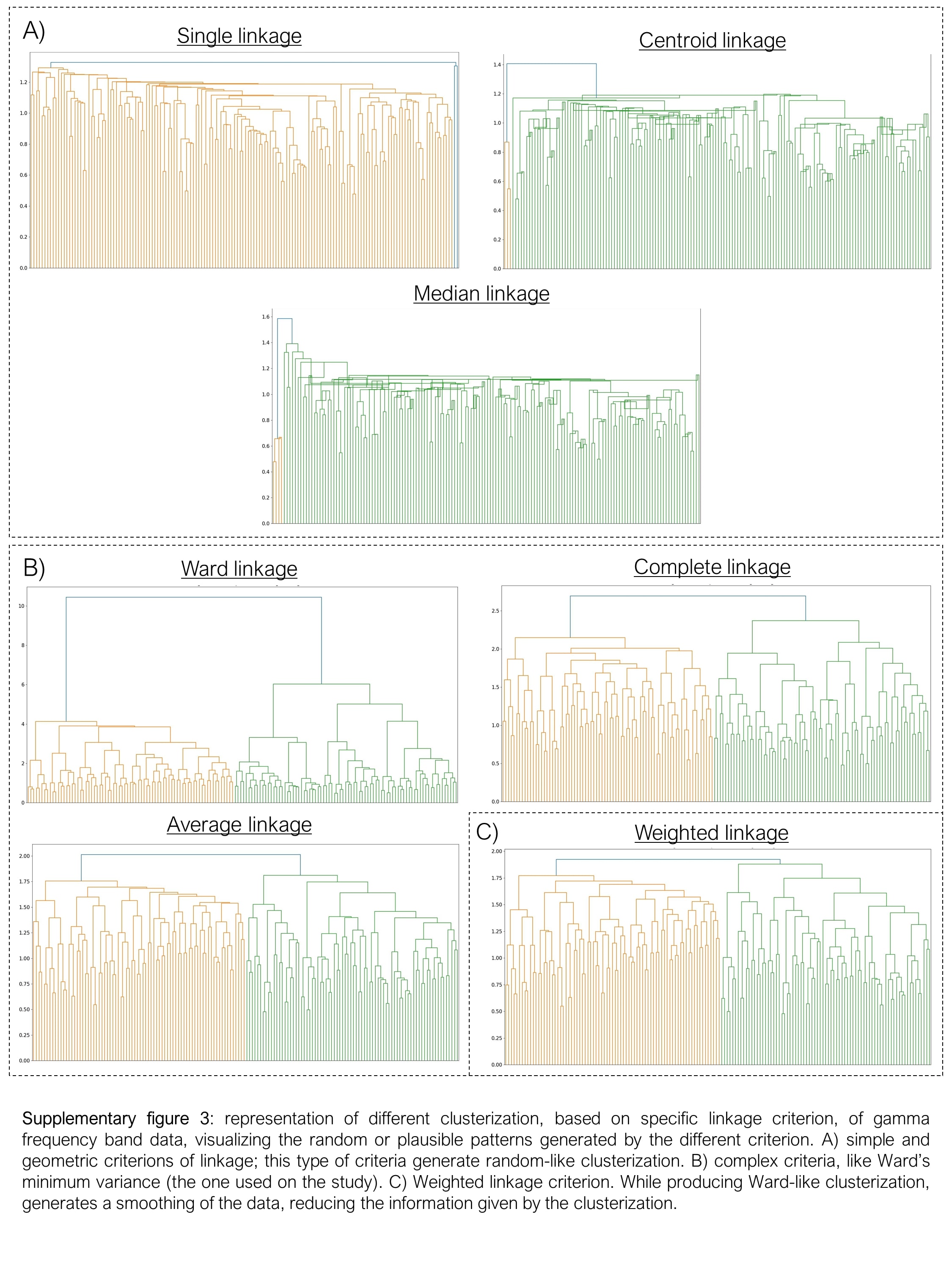

### Supplementary Table 1

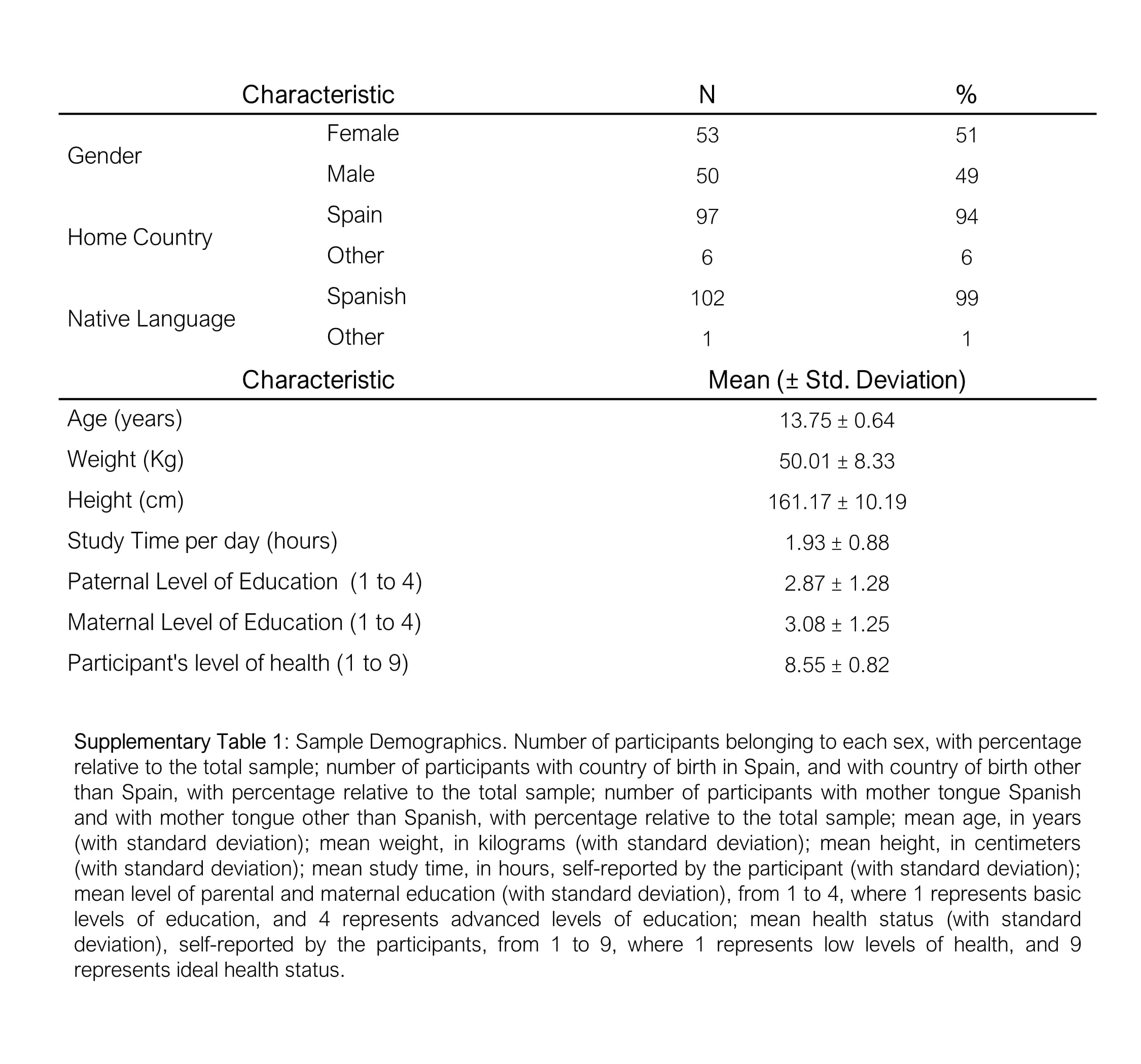

### Supplementary Table 2

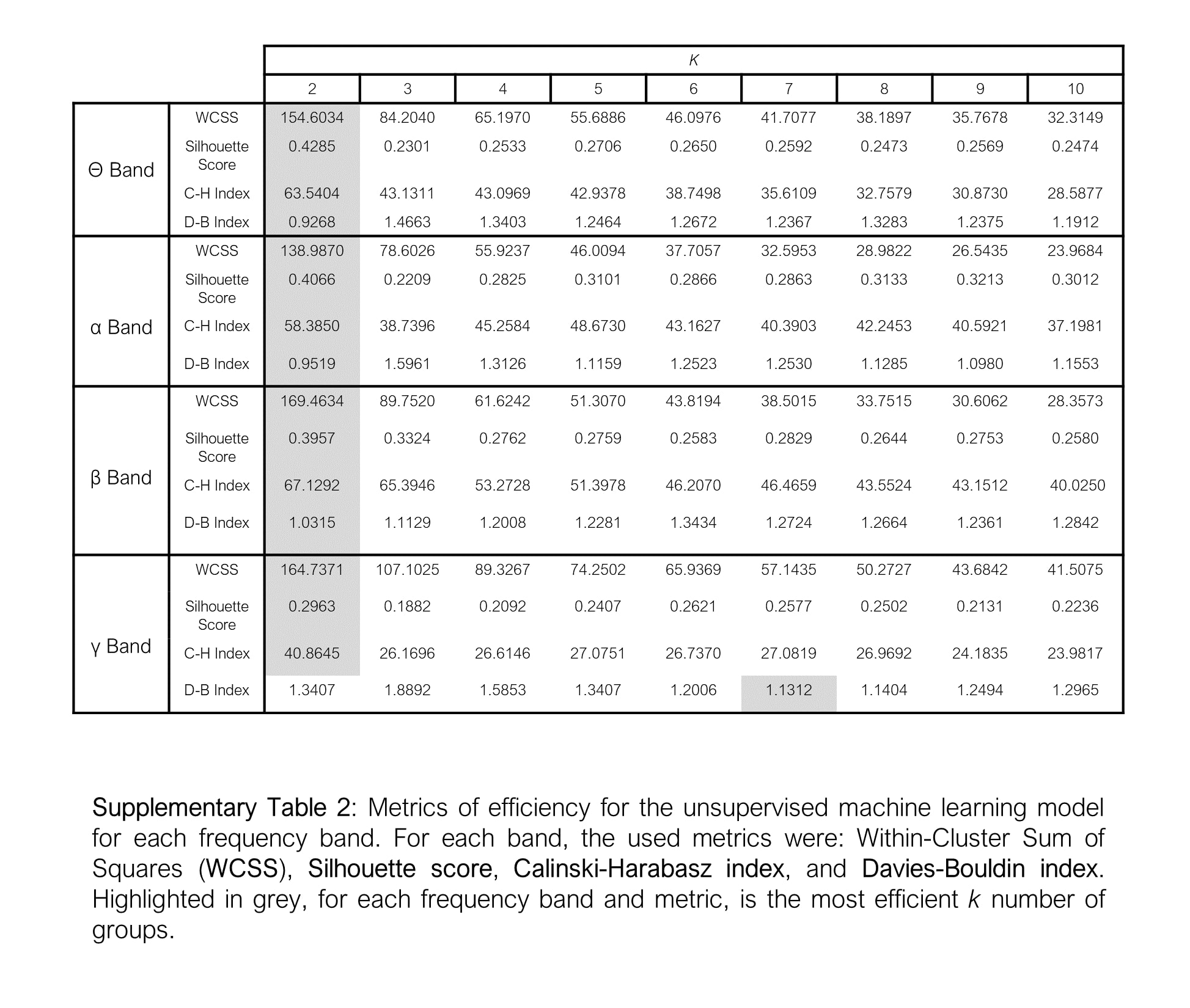
